# Investigation of the cellular reprogramming phenomenon referred to as stimulus-triggered acquisition of pluripotency (STAP)

**DOI:** 10.1101/027730

**Authors:** Hitoshi Niwa

## Abstract

In 2014, it was reported that strong external stimuli, such as a transient low-pH stressor, was capable of inducing the reprogramming of mammalian somatic cells, resulting in the generation of pluripotent cells (Obokata et al. 2014a, b). This cellular reprograming event was designated ‘stimulus-triggered acquisition of pluripotency’ (STAP) by the authors of these reports. However, after multiple instances of scientific misconduct in the handling and presentation of the data were brought to light, both reports were retracted. To investigate the actual scientific significance of the purported STAP phenomenon, we sought to repeat the original experiments based on the methods presented in the retracted manuscripts and other relevant information. As a result, we have confirmed that the STAP phenomenon as described in the original studies is not reproducible.

## Introduction

Cellular reprograming is a biological event in which a differentiated metazoan cell is induced to revert to a state functionally resembling that of cells at earlier developmental stages. Full reprograming of somatic cells results in the acquisition of the ability to give rise to an entire organism, or totipotency; this can be achieved by somatic cell nuclear transfer (Gurdon 1962). Pluripotency in contrast is the ability of a cell to differentiate into all somatic cell lineages. It has been shown that the artificial expression of pluripotency-associated transcription factors results in reprogramming of somatic cells to a state of pluripotency, such cells are referred to as as induced pluripotent stem (iPS) cells (Takahashi & Yamanaka 2006).

Mouse pluripotent stem cells share common features. Authentic pluripotent stem cells are embryonic stem (ES) cells derived from pre-implantation embryos (Evans & Kaufman 1981; Martin 1981). Under optimized culture conditions, these maintain self-renewal by giving rise to pluripotent daughter cells via cell division. Leukemia inhibitory factor (LIF) is a well-known factor sufficient to maintain the pluripotency of mouse pluripotent stem cells *in vitro* (Smith *et al.* 1988). Such cells express a unique set of genes associated with pluripotency, such as a transcription factor *Oct3/4* (Okamoto *et al.* 1990), and contribute to embryo development when transferred into pre-implantation embryos, resulting in the formation of germline chimeras (Bradley *et al.* 1984). These properties are shared by iPS cells derived from somatic cells (Takahashi & Yamanaka 2006). Therefore, acquisition of pluripotency by somatic cells via reprograming is typically assessed based on such criteria.

In 2014, it was reported that sublethal external stimuli, such as exposure to a transient low-pH stressor, reprogrammed mammalian somatic cells, resulting in the generation of pluripotent cells (Obokata *et al.* 2014a; Obokata *et al.* 2014d). In these reports, this cellular reprograming event was designated ‘stimulus-triggered acquisition of pluripotency’ (STAP). The reports also described how the primary pluripotent cells, STAP cells, were able to give rise to two types of pluripotent stem cells in a culture-condition-dependent manner. However, both reports were subsequently retracted due to multiple intsances of scientific misconduct (Obokata *et al.* 2014b; Obokata *et al.* 2014c). To investigate the scientific significance of the STAP phenomenon, we repeated the reported experiments based on the methods presented in the retracted manuscripts and other relevant information subsequently obtained. We examined the expression of pluripotency-associated genes in cell aggregates obtained in cultures of somatic cells treated with transient low-pH, and ability of such cell aggregates to contribute to chimeric embryos after injection into pre-implantation embryos. The results of this reevaluation indicate that the previously reported STAP phenomenon is not reproducible.

## Results

### Transient low-PH treatment enhances formation of characteristic cell aggregates

In the original report, a transient low-pH stress induced by addition of hydrochloric acid (HCl) caused massive cell death of dissociated somatic cells around 1–2 days after treatment (Figure 1d of Obokata *et al.* 2014a); the surviving cells formed aggregates (Figure 1b of Obokata *et al.* 2014a). In the present study, we examined the effect of HCl treatment on dissociated cells derived from spleen, liver and heart of 4–9-day-old mice. The amount of the diluted HCl solution to achieve optimized low-pH condition was adjusted (to around pH=5.7, Fig. 1a) as indicated in the previous manuscript (Figure S1a of Obokata *et al.* 2014a) and experiments were repeated several times. However, although massive cell death was observed at two days after treatment, aggregate formation was rarely observed in any cell type (Fig. 1b). Occasional formation of aggregates was also observed in the culture of non-treated cells, suggesting that low-pH treatment does not enhance the formation of cell aggregates.

**Figure 1.**
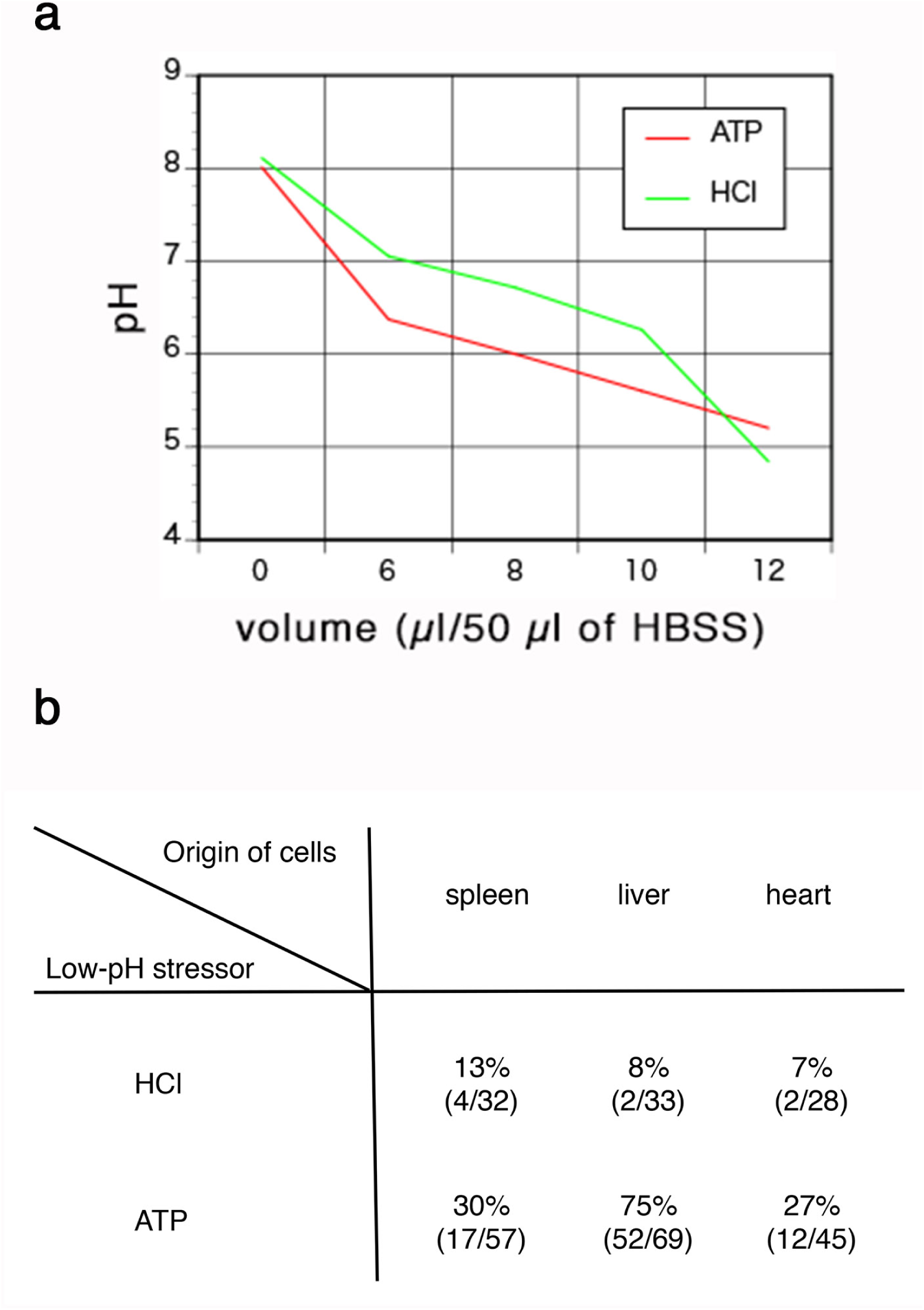
Optimization of the condition for low-pH treatment. **a.** Titration of HCl and ATP to achieve optimal low-pH condition of cell suspension. Indicated volumes of the diluted HCl or ATP solution was added to 500 *µ*l of HBSS containing 7 10^5^ liver cells and pH was measured. **b.** Frequency of formation of cell aggregates from the cells prepared from various tissues after low-pH treatment. The numbers of the total experimental trials and the trials with formation of cell aggregates at each combination of cell types and low-pH stressors are indicated.

Next we examined the effect of adenosine triphosphate (ATP) as a transient low-pH stressor based on personal communication with the authors of the original study. The amount of the diluted ATP solution to achieve optimized low-pH (~5.7) was adjusted (Fig. 1a) and experiments were repeated several times. Massive cell death was again observed at two days after treatment (Fig. 2a); however, we found that liver cells reproducibly gave rise to cell aggregates morphologically similar to those shown in the previous report, whereas spleen and heart cells only occasionally formed similar cell aggregates (Fig. 1b). The efficiency of aggregate formation was clearly higher for ATP-treated cells than for HCl-treated or non-treated cells, especially in the case of liver cells.

**Figure 2.**
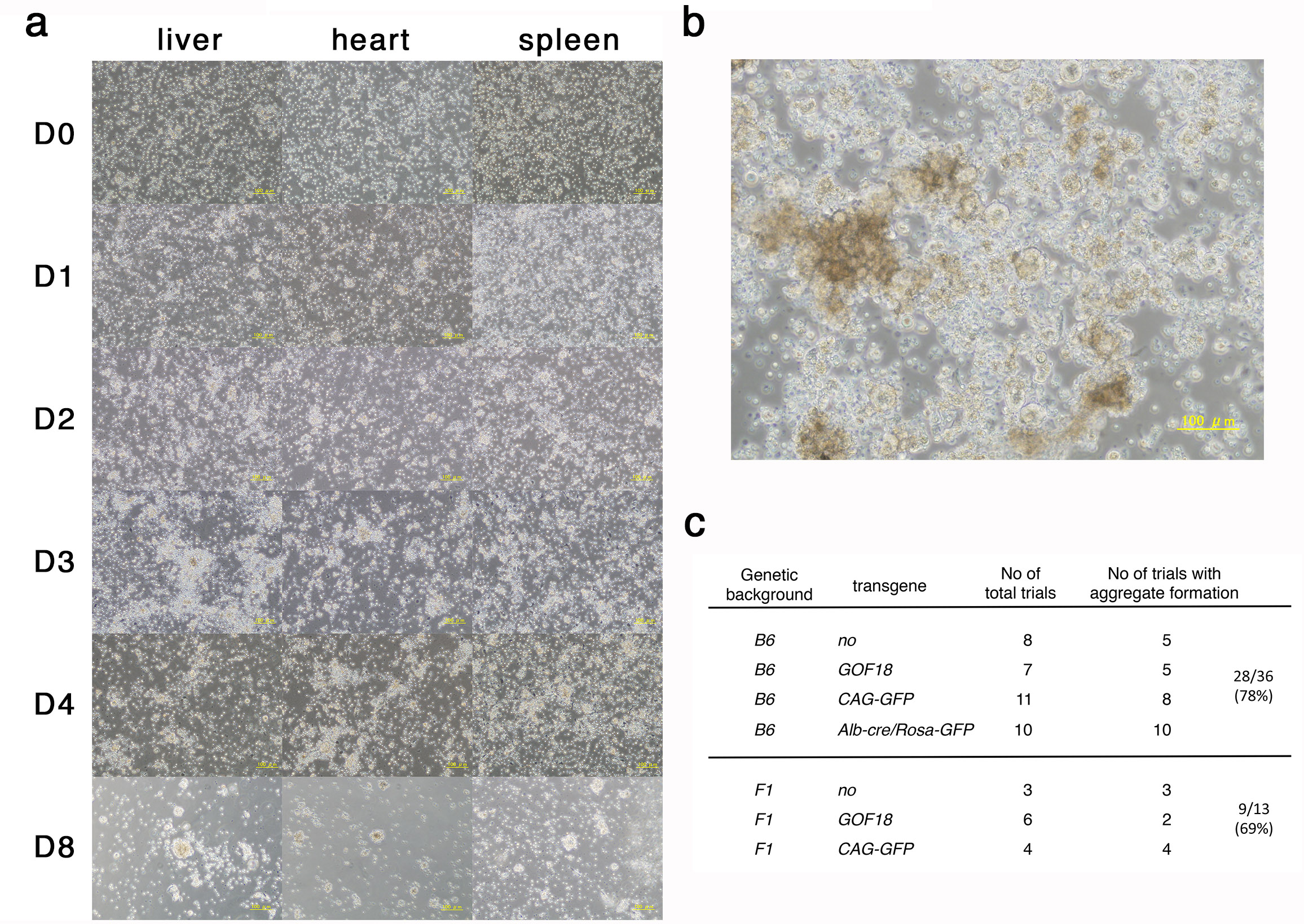
Formation of cell aggregates from low-pH treated cells. **a.** Time course of the cultures of liver, heart and spleen cells treated with ATP. The cells were prepared from 5-days old of C57BL6 mice carrying CAG-GFP. Scale bar = 100 *µ*m. **b.** Cell aggregates derived from liver cells treated with ATP with the culture for 7 days. Liver cells were prepared from 4-days old of C57BL6/129 F1 mice. Scale bar = 100 *µ*m. **c.** Frequency of formation of cell aggregates from liver cells with different genetic backgrounds. B6; C57BL6, F1; C57BL6/129 or 129/C57BL6. The numbers of the total experimental trials and the trials with formation of cell aggregates at each combination of cell types and genetic backgrounds are indicated.

In the case of liver cells, when 5 10^5^ cells were seeded in a well of 12 well plate, 20–30 aggregates were observed after seven days on average (Fig. 2b). Addition of fibroblast growth factor (Fgf)-2 based on personal communication with the authors slightly enhanced cell agregate formation. Since the culture medium contains leukemia inhibitory factor (LIF), which shows differential action on ES cells derived from different genetic backgrounds (Ohtsuka & Niwa 2015), we suspected that the genetic background might affect aggregate formation. However, as shown in Fig. 2c, although it was known that the *129* background confers a dominant effect in obligating the LIF signal input to maintain pluripotency (Ohtsuka & Niwa 2015), there was no difference between *C57BL6* and *C57BL6 x 129 F1* (either *C57BL6/129* or *129/C57BL6*) in the observed frequency of aggregate formation in the present study.

### Induced cell aggregates show poor induction of pluripotency-associated markers

To test the induction of pluripotency markers in cell aggregates obtained from ATP-treated liver cells, we assessed the expression of pluripotency-associated genes. *Oct3/4* is a well-defined marker of pluripotent stem cells. Using a primer pair to detect *Oct3/4* transcript from the *Pou5f1* allele, but not pseudo-genes (Mizuno & Kosaka 2008), we did not find a detectable level (above 0.1% of the expression level in mouse ES cells, relative to the expression levels of *Gapdh*) of the transcript by quantitative polymerase chain reaction (Q-PCR) using a total RNA sample prepared from all cells in the culture (Fig. 3a), indicating that extremely few or no cells expressing *Oct3/4* were present. Interestingly, expression of *Gfp* from the *Oct3/4-GFP* transgene (*GOF18*) (Yeom *et al.* 1996) was detected in liver cells cultured for seven days irrespective of ATP treatment, suggesting leaky expression of this transgene.

**Figure 3.**
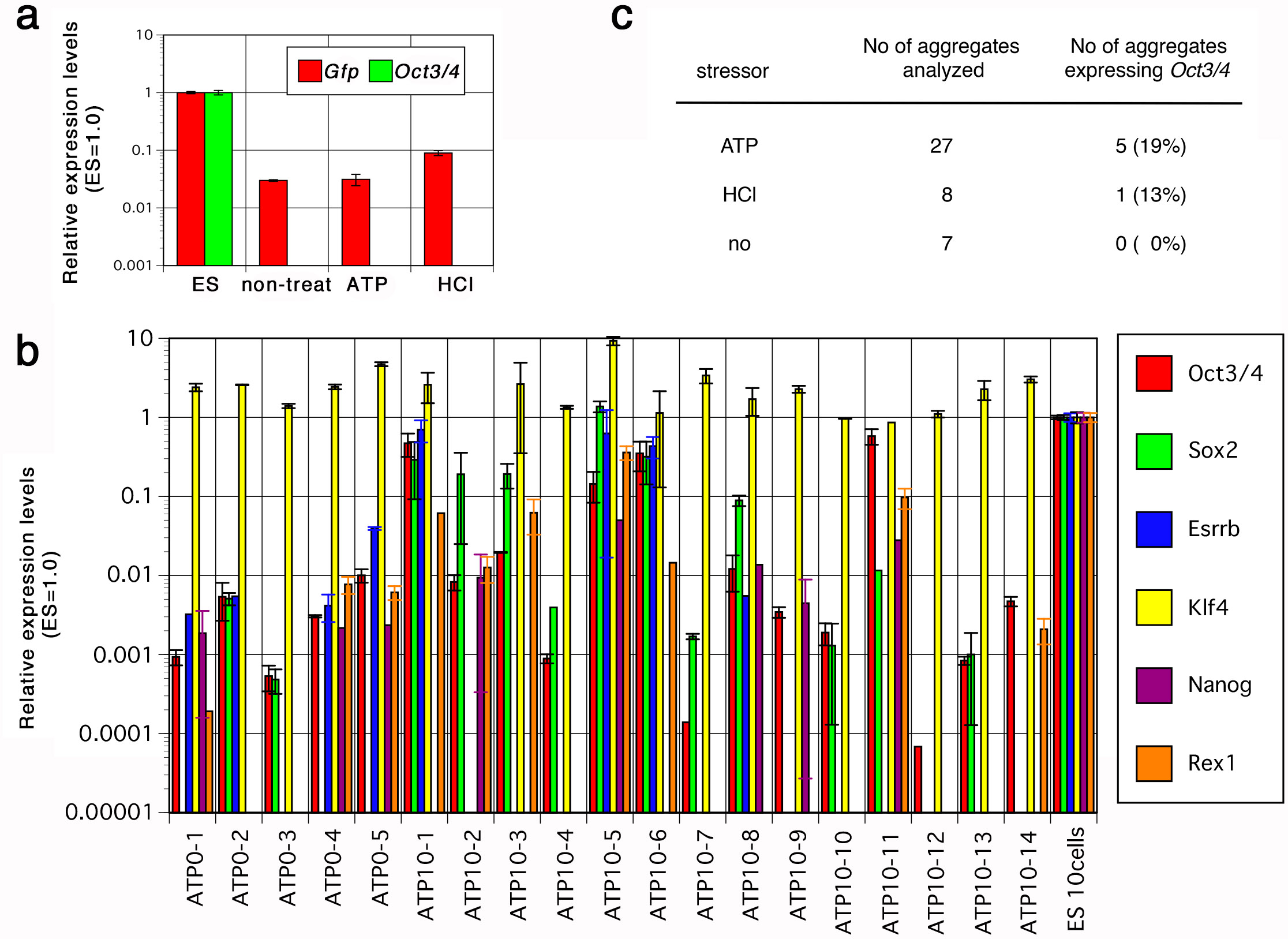
Q-PCR analysis for the expression of pluripotency markers in induced cell aggregates. **a.** Q-PCR analysis of the low-pH treated liver cells cultured for 7 days. Liver cells were prepared from 7-day old *GOF* mice and treated with either ATP or HCl, or without stressor. RNA samples were prepared from all cells in the wells at day 7 of culture and the relative expression levels of *Gfp* (derived from *GOF Tg*) and *Oct3/4* (derived from the endogenous *Pou5f1* allele) to *Gapdh* were indicated with standard deviation. The expression levels in control ES cells carrying *CAG-GFP Tg* were set at 1.0. **b.** Q-PCR analysis of the single cell aggregates derived from the ATP-treated or non-treated liver cells cultured for seven days. The liver cells were prepared from 4-days old of *C57BL6/129* mice and the single cell aggregates were separately treated for quantification of gene expression. The relative expression levels of pluripotency-associated genes to *Gnb2l1* were indicated with standard deviation. The expression levels in 10 control ES cells were set at 1.0. **c.** Frequency of cell aggregates showing significant levels of *Oct3/4* expression. The relative expression levels of *Oct3/4* in single cell aggregates derived from liver cells were measured as b and the frequency of the cell aggregates with significant levels of *Oct3/4* expression (over 0.001 of relative expression) is indicated.

We next performed Q-PCR on individual cell aggregates isolated from culture. Aggregates were selected and RNA samples were prepared separately. These RNAs were reverse-transcribed and QPCR was performed. We found that some aggregates expressed a significant amount—more than 10% of the expression level in ES cells—of pluripotency-associated genes, including *Oct3/4* (Fig. 3b). *Klf4* expression was detected in all samples, which may reflect its expression in liver cells and thus serve as a positive control of this assay. Of cell aggregates derived from liver cells treated with ATP, 19% expressed a significant amount of *Oct3/4* (Fig. 3c). These data suggest that some proportion of cells in the aggregates express pluripotency-associated genes at comparable levels to those of ES cells.

To examine the proportion of the cells expressing Oct3/4 in the aggregates, we next applied immuno-staining using a specific antibody against Oct3/4 we raised and assessed previously (Niwa *et al.* 2005). Cell aggregates derived from low-PH treated liver cells were fixed, stained by anti-Oct3/4 antibody, and observed using confocal microscopy. We stained morula-stage mouse embryos as positive controls. By comparison with these positive controls, we found that some of the cell aggregates contained cells expressing Oct3/4 at comparable levels (Fig. 4a). In the case of cell aggregates derived from liver cells treated by ATP, 20% of cell aggregates contained Oct3/4-positive cells (Fig. 4b), which is consistent to the proportion of cell aggregates expressing significant amounts of *Oct3/4* detected by QPCR (Fig. 3c). In contrast, cell aggregates derived from liver cells treated by HCl included Oct3/4-positive cells at a frequency comparable to that of non-treated cells. The presence of Oct3/4-positive cells in the cell aggregates occasionally found in cultures of non-treated liver cells suggests that such cells are derived from Oct3/4-positive cells present in the liver cell population, that *in vitro* culture may itself be a source of the stress to the cells, or that the immuno-staining technique may produce some non-specific signal. In cell aggregates derived from ATP-treated liver cells, the Oct3/4-positive cells were typically positioned in the center of the cell aggregates and exhibit large nuclei, and were surrounded by Oct3/4-negative cells with small nuclei at the peripheries of the cell aggregates (Fig. 4c).

**Figure 4.**
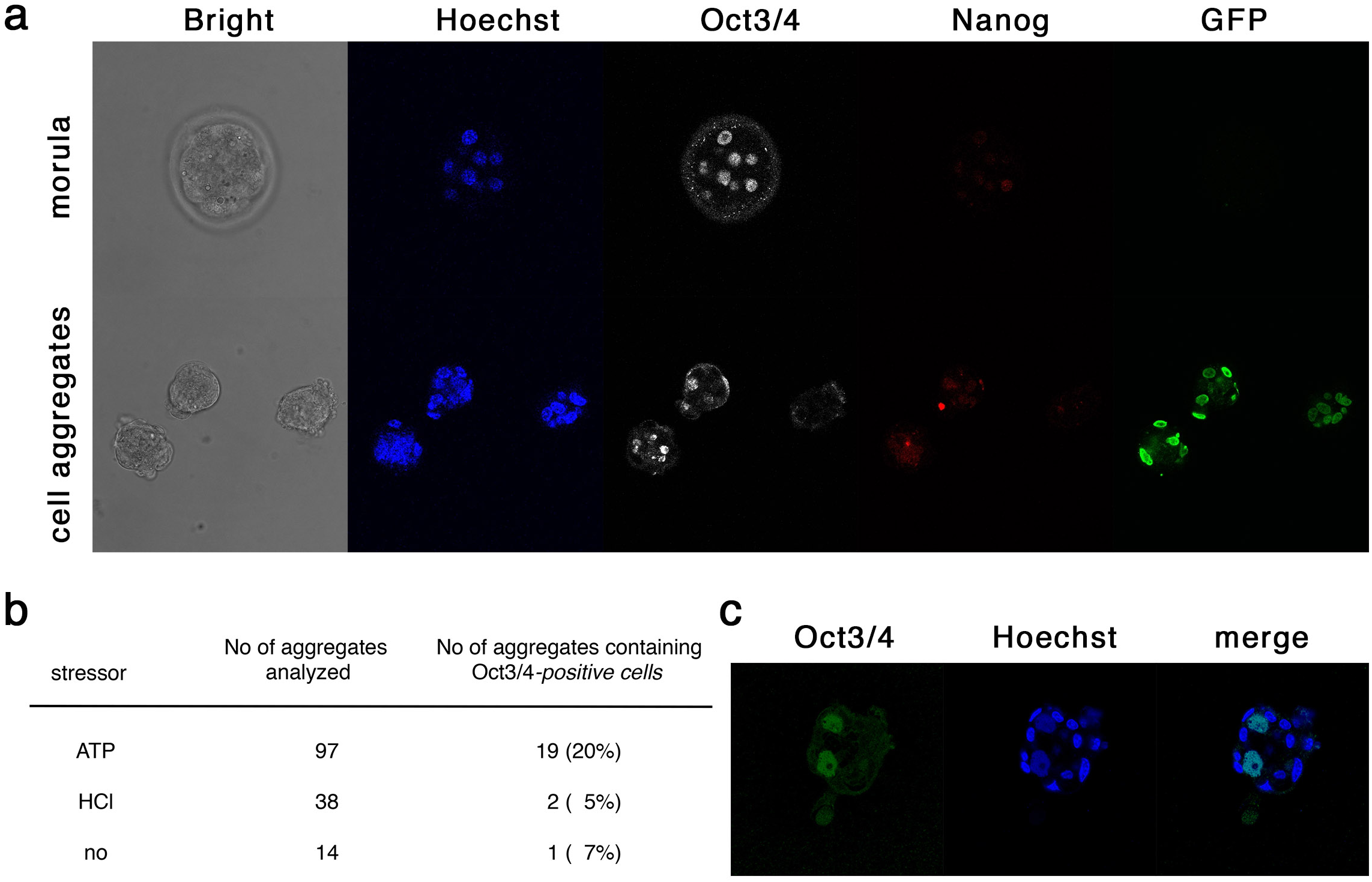
Immuno-staining of cell aggregates derived from low-pH treated liver cells. **a.** Immunostaining of morula-stage embryos and cell aggregates for Oct3/4 and Nanog. Both samples were treated in parallel and confocal microscopic images were captured with the same exposure time. The embryo is wild-type *C57BL6* whereas the cell aggregates were derived from 8-days old of *Alb-cre/Rosa-GFP Tg* mice. **b.** Frequency of cell aggregates carrying Oct3/4-positive cells. The numbers of the immune-stained cell aggregates derived from liver cells and that of carrying Oct3/4-positive cells are indicated for each stressor treatment. **c.** Immuno-staining image of cell aggregate for Oct3/4 derived from liver cells prepared from 4-days old C57BL6/129 mice.

### *Oct3/4-GFP* transgene expression not detected in low-pH treated cells

In the original report, the authors used transgenic reporter gene expression as a marker of pluripotency (Obokata *et al.* 2014a). This reporter consisted of the transcriptional regulatory element of *Oct3/4* and the fluorescent marker *GFP*, designated *GOF*, which is silent in somatic cells and activated in pluripotent cells (Yeom *et al.* 1996). When we used the same transgenic mouse line as a source of dissociated cells, we found that they began to acquire strong auto-fluorescence in culture after ATP treatment. By observation with fluorescent microscopy, most of the aggregates showed both green and red fluorescence, a sign of auto-fluorescence (Fig. 5a) although this may also include the fluorescent signal from the *GOF* transgene, since we detected *Gfp* mRNA by QPCR in these cells after culture *in vitro* for seven days (Fig. 3a).

**Figure 5.**
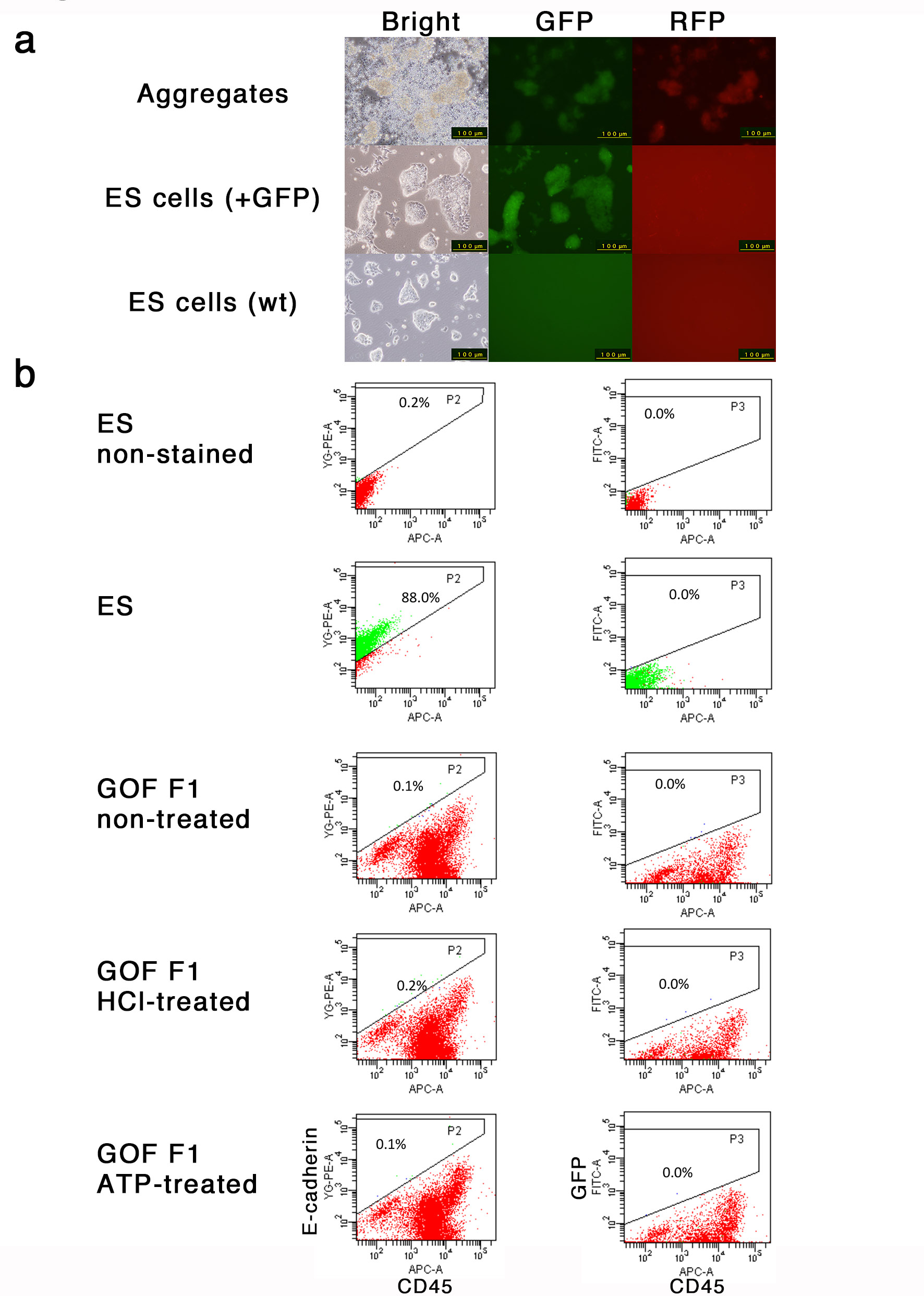
Analyses of fluorescent signals from *GOF* transgene. **a.** Fluorescent microscopic analysis of cell aggregates derived from *GOF Tg* mice. The cell aggregates were derived from liver cells of 6-days old of GOF Tg mice. Fluorescent images with the filter sets for detection of GFP and RFP signals are shown. Images of ES cells carrying *CAG-GFP* captured with the same conditions are shown as a control. **b.** FACS analysis of the low-pH treated spleen cells derived from *GOF Tg* mice. The spleen cells were isolated from 7-day-old *GOF Tg* mice and prepared with Lympholyte followed by treatment with the indicated stressors. After the culture for seven days, the cells were dissociated, stained with anti-E-cadherin with PE and anti-CD45 with APC, and analyzed by FACS. Wild-type ES cells were used as a positive control for E-cadherin staining and a negative control for CD45-staining as well as GFP fluorescence.

Specific detection of GFP fluorescence by fluorescence-activated cell sorting (FACS) was also applied. In spleen cells collected using Lympholyte, CD45-positive/E-cadherin-negative blood cells were enriched. The reprogramming of such cells to a state of pluripotency can be monitored by their conversion to CD45-negative/E-cadherin-positive cells and acquisition of GFP expression from the *GOF* transgene. However, although we again observed increased auto-fluorescence and some reduction of CD45 expression, neither a specific signal of GFP fluorescence nor an increase of E-cadherin expression was observed in the low-pH treated cells (Fig. 5b). Given these findings, we suggest that the *GOF* fluorescence marker is unsuitable for use as a marker of up-regulation of *Oct3/4* under these experimental conditions, and that there was no evident sign of reprogramming in low-pH treated spleen cells.

### Induced cell aggregates do not contribute to chimeric embryos after injection into pre-implantation embryos

In the original report, the authors showed that the cell aggregates obtained by the culture of low-pH treated cells contribute to chimeras when the cell aggregates were chosen by their morphologies under the microscopic observation, manually dissected and injected into blastocysts (Figure 4a of (Obokata *et al.* 2014a)). The frequency of obtaining chimeric mice from injected blastocysts reached 24% (64 chimeric mice from 264 injected blastocysts; Figure S7b of (Obokata *et al.* 2014a)). We prepared cell aggregates from the liver cells dissociated from the livers of transgenic mice carrying *CAG-EGFP* (Okabe *et al.* 1997) or selected cell aggregates expressing GFP prepared from the liver cells derived from *Alb-cre: Rosa-GFP* double transgenic mice (Abe *et al.* 2011; Postic *et al.* 1999) (Fig. 6, top) and repeated injections of these into morula and blastocysts eight times. However, we found no chimeric embryos carrying GFP-positive cells among 117 embryos derived from 244 injected embryos based on observation by fluorescent microscopy (Fig. 6, bottom). These data strongly suggest that the acquisition of pluripotency rarely occurs in cell aggregates derived from low-pH treated liver cells, if at all.

**Figure 6.**
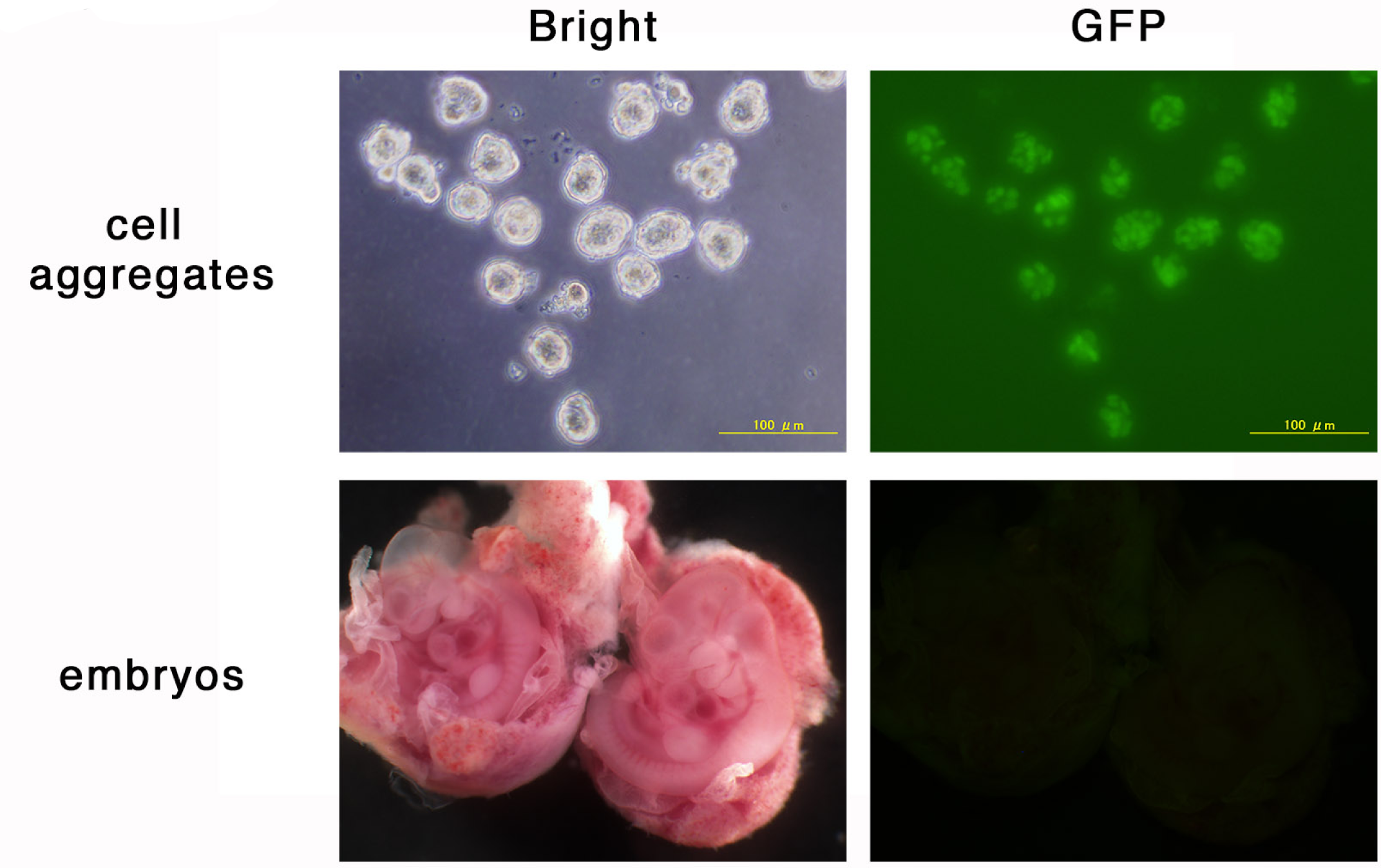
Chimera assay of cell aggregates. Cell aggregates derived from liver cells prepared from 8-day-old *Alb-cre:Rosa-GFP Tg* (upper panels) were injected into morula-stage embryos. The manipulated embryos were transferred into the uterus and the embryos were recovered at E10.5. The dissected embryos were observed under fluorescent microscopy for the contribution of GFP-positive cells.

### Induced cell aggregates do not give rise to stem cell lines

The original studies reported that two different types of stem cell lines could be established from cell aggregates obtained by the culture of low-pH treated cells: ES-like ‘STAP stem cells’ (Figure 5 of (Obokata *et al.* 2014a)) and trophoblast stem (TS)-like ‘FGF-induced (FI) stem cells’ (Figure 2 of (Obokata *et al.* 2014b)). To reevaluate these reports, we transferred cell aggregates derived from liver cells from various genetic backgrounds into the culture conditions for derivation of either ES-like or TS-like stem cells. In the case of the culture for ES-like stem cells in serum-free culture containing knockout serum replacement (KSR), adrenocorticotropic hormone (ACTH) and LIF (Ogawa *et al.* 2004), most of the cell aggregates died without outgrowth, which may attributable to the absence of serum, while a small number aggregates gave rise to colonies containing small cells with large nuclei, resembling the morphology of embryonic stem cells. However, most of these cells ceased proliferation at day 7 and gradually regressed. Significant proliferation after day 7 was observed in only three of 492 cell aggregates and none of these gave rise to cell lines (Fig. 7a). In the case of the culture for TS-like stem cells containing FGF-4 and heparin (Tanaka *et al.* 1998), many clumps showed outgrowth of fibroblastic cells, which may be due to the presence of FGF2 in the medium. Few of these (22 of 391 cell aggregates) gave rise to colonies of small stem cell-like cells and one of them could be passaged three times (Fig. 7b). However, all of them ultimately regressed without giving rise to cell lines. These data showed that we are unable to derive stem cell lines from aggregates derived from low-pH treated liver cells.

**Figure 7.**
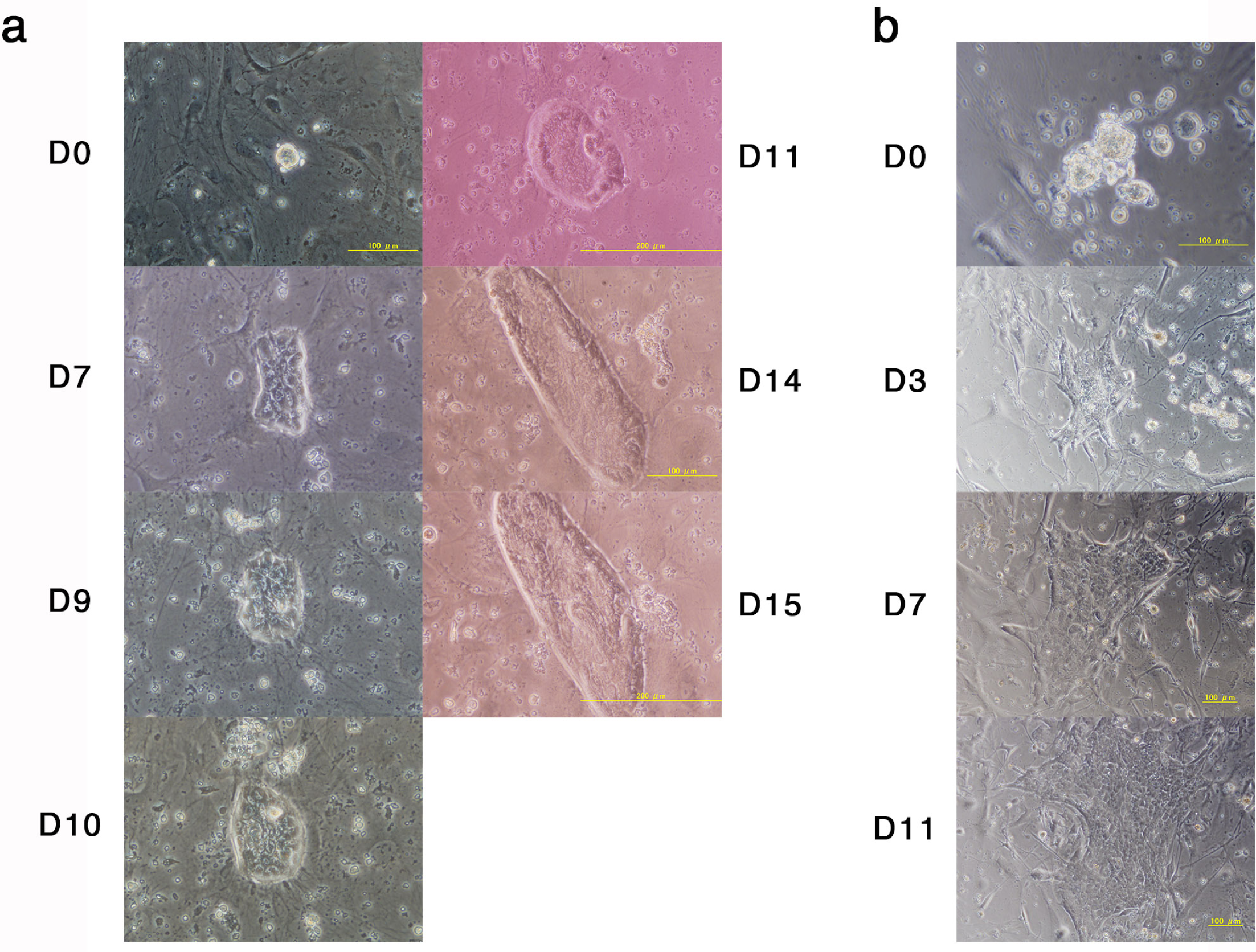
Culture of cell aggregates *in vitro*. **a.** The outgrowth culture of cell aggregate derived from liver cells. Liver cells were prepared from 7-days old of *C57BL6 CAG-GFP Tg*, treated with ATP and cultured for six days. Single cell aggregates were isolated and cultured on MEF feeder cells with medium containing KSR, ACTH and LIF adapted to the culture of ES cells. The cells continue to grow for 15 days but did not give secondary colony after passage. **b.** Outgrowth culture of cell aggregates derived from liver cells. Liver cells were prepared from 4-day-old *C57BL6/129* mice, treated with ATP, and cultured for six days. The cell aggregates were isolated and cultured on MEF feeder cells with medium containing FGF4 and heparin adapted to the culture of TS cells. The cells continued to grow for 11 days, but did not give rise to secondary colonies after passage.

## Discussion

In the present study, we investigated the properties of cell aggregates obtained by culture of liver cells transiently treated with low-pH stimulus. Interestingly, few cells in a subset of cell aggregates expressed significant amounts of the pluripotency marker *Oct3/4*, but the frequency was very low; 5 × 10^5^ liver cells yielded only ~30 cell aggregates, in which about 20% of the cell aggregates contain 1–2 Oct3/4 positive cells, indicating a frequency per seeded liver cell of 0.0012–0.0024%. Moreover, the pluripotency of such cells was not confirmed by chimera formation assay and they did not give rise to any stem cell lines. We thus conclude that such cell aggregates do not fulfill the definition for STAP cells proposed in the original studies. Moreover, since the frequency of Oct3/4-positive cells in the cell aggregates was quite low, it was impossible to determine whether they were selected from the original population or induced in culture, again highlighting the lack of clear evidence for the existence of the reported STAP phenomenon.

## Materials and methods

### Animals

*C57BL/6NJcl* (CLEA Japan) and *129X1/SvJJmsSlc* (Japan SLC) mice were purchased from suppliers. *C57BL/6-Tg(CAG-EGFP)C14-Y01-FM131Osb* transgenic mouse (*CAG-GFP Tg*) line was provided by Research Institute for Microbial Disease, Osaka University (Okabe *et al.* 1997). *C57BL/6J-Tg(GOFGFP)11Imeg* transgenic mouse (*GOF-Tg*) line was obtained from RIKEN Bio-Resource Center (RBRC00771) (Ohbo *et al.* 2003). *B6.Cg-Tg(Alb-cre)21Mgn/J* transgenic mouse (*Alb-cre Tg*) line was supplied by Jackson Laboratory (Postic *et al.* 1999). *R26R-H2B-EGFP* transgenic mouse (*Rosa-GFP*) line was generated by Laboratory for Animal Resources and Genetic Engineering (LARGE), RIKEN CDB (Abe *et al.* 2011).

### Isolation of cells from mice

4–9-day-old mice were euthanized using carbon dioxide and then sterilized with 70% ethanol. For the isolation of spleen cells, excised spleen was minced with scissors and the tissue fragments were dissociated in phosphate buffered serine (PBS) by pipetting. The cell suspension was strained through a cell strainer followed by the collection of cells by centrifugation at 1,000 rpm for 5 min. The collected cells were re-suspended in 5 ml of Dulbecco’s Modified Eagle medium (DMEM; Life Technologies) and added to the same volume of Lympholyte^®^ (Cedarlane), and then centrifuged at 1,000 g for 20 min. The lymphocyte layer was isolated and washed with PBS to obtain single cell suspension. For the isolation of liver cells, excised liver was minced with scissors and the tissue fragments were dissociated by incubation in Type I collagenase (Worthington Biochemical) solution (0.5 mg/ml in Hanks Balanced Salt Solution (HBSS, no calcium, no magnesium; Life Technologies)). Next, the cell suspension was strained through a cell strainer followed by the collection of cells by centrifugation at 1,000 rpm for 5 min. For the isolation of heart cells, the excised heart was minced with scissors and the tissue fragments were dissociated by incubation in Type II collagenase (Worthington Biochemical) solution (0.5 mg/ml in HBSS). The cell suspension was strained through a cell strainer followed by the collection of cells by centrifugation at 1,000 rpm for 5 min.

### Low-pH treatment and culture of cell aggregates

Diluted HCl solution was prepared with 10 *µ*l of 35% HCl (Nakarai) in 590 *µ*l HBSS. Diluted ATP solution was prepared with ATP (Sigma) in distilled water at 200 mM. Titration of pH with various amount of diluted HCl or ATP was performed with 500 *µ*l of HBSS containing 7 10^5^ liver cells. As a routine method, 10 *µ*l of either diluted HCl or ATP solution was added into 500 *µ*l of cell suspension containing 5 10^5^ cells in HBSS followed by incubation for 25 min at 37˚C, and then centrifuged at 1,000 rpm at room temperature for 5 min. After the supernatant was removed, precipitated cells were re-suspended and plated onto either adhesive or non-adhesive plates at cell density of 1–5 10^5^ cells per well in 1 ml of the culture medium. The culture medium consists of DMEM/HamF12 (Life Technologies) supplemented with 1,000 U/ml of mouse LIF (home-made) and 2% of B27^®^ Supplement (Life Technologies). Optionally, recombinant human Fgf2 (Wako) was added at final concentration of 10 ng/ml.

### QPCR

To quantify the levels of mRNA transcripts, total RNA was prepared by TRIzol^®^ (Life Technologies). cDNA were synthesized from 1 *µ*g of total RNA using SuperScript^®^ III (Life Technologies), and quantified by real-time PCR using a CFX384 system (BioRad). Utilized primers were listed on Table 3. All samples were tested in triplicate, and the mean relative amounts of each transcript were calculated by normalization to an endogenous control *Gapdh*.

Each cell aggregate was washed with PBS and transferred in 2 *µ*l of PBS into 8 *µ*l RealTime ready Cell Lysis Buffer (Roche) supplied with NP-40, RNAsin and RNase inhibitor. Then 3 *µ*l of cell lysis solution was mixed with 1.5 *µ*l of DNaseI solution (0.2 U/*µ*l) to degradate genomic DNA followed by addition of 1.5 *µ*l of 8 mM EDTA solution to stop the reaction. For reverse transcription of RNA, 3 *µ*l of pre-mixture of SuperScript^®^ VILO reverse transcriptase (Life Technologies) was added into 6 *µ*l of DNaseI-treated cell lysate and incubated at 42˚C for 1 hour. The reverse-transcribed product was pre-amplified with Plutinum multiplex PCR master mix using pooled primer mixture using the reaction cycle (95˚C for 30 sec; 60˚C for 90 sec; 72˚C for 60 sec) for 14 cycles. The mixture was treated with Exonuclease I to remove the primers for pre-amplification, and quantitative PCR was performed with the primer pairs specific for each gene using Quantitest SYBR Green PCR mix (Qiagen) in BioRad CFX384 Real-Time System (Bio-Rad). Utilized primers were listed on Table 4. All samples were tested in triplicate, and the mean relative amounts of each transcript were calculated by normalization to an endogenous control *Gapdh or Gnb2l1*.

### Immunostaining

Cells were fixed by 4% paraformaldehyde in PBS for 30 min at 4°C and then permeabilized by 0.1% Triton X-100 in PBS for 15 minutes at room temperature (RT). After brief washing with PBS followed by blocking with PBS containing 2% FCS, the cells were incubated with the following primary antibodies: anti-Oct3/4 rabbit antiserum (Niwa *et al.* 2005) and anti-Nanog rat monoclonal antibody (R&D) for overnight at 4°C. After washing with PBS, the cells were incubated with Alexa Fluor 488-or 633-conjugated donkey antibodies (Invitrogen) were used in a proper combination of species specificity as indicated in Figure legends. Fluorescent images were captured with an IX51 microscope with DP70 digital camera (Olympus) or a Leica SP8 confocal microscope (Leica).

### FACS

For flow cytometric analyses, cell aggregates were harvested, washed by PBS, and incubated with TrypLE^TM^ Select (Life Technologies) for 5 min. After dilution with culture medium, aggregates were dissociated into single cells by gentle pipetting. Cells adhered to the culture substrate were also harvested following a standard method. These cells were mixed and collected as pellets by a centrifugation. For quantification of GFP-positive population, dissociated cells were re-suspended in 500 *µ*l HBSS containing 1 *µ*l of DRAQ7, a cell-nonpermeable DNA dye (for the detection of dead cells; Cell Signaling). When combined with a staining for CD45 antigen, cell pellets were suspended with 50 *µ*l HBSS containing 10 *µ*l of APC-conjugated rat anti-CD45 antibody (BD Pharmingen), and incubated for 30 min on ice. For co-staining with CD45/E-cadherin antibodies, cell pellets were suspended in culture medium, and then incubated for 30 min in CO_2_ incubator. The cells were harvested and suspended with 50 *µ*l HBSS containing 5 *µ*l biotin-labeled rat anti-E-cadherin antibody (ECCD2). After incubation for 30 min on ice, the stained cells were once washed by HBSS, re-suspended with 50 *µ*l HBSS containing 1 *µ*l PE-conjugated streptavidin (Life Technologies) and 10 *µ*l APC-conjugated anti-CD45 antibody, and further incubated for 30 min on ice. These stained cells were once washed by HBSS and suspended with 500 *µ*l HBSS.

After the cell suspension was passed through a filter mesh, the cells were analyzed using a FACSAria IIIu cell sorter (Becton Dickinson).

### Injection into pre-implantation embryos

Cell aggregates were cut into smaller pieces using a laser (XYClone, Nikko Hansen & Co., Ltd) or shaped glass capillaries, which were then microinjected into 8-cell or blastocyst stage embryos from *ICR* mice (Charles River Laboratories Japan, Inc.). Injected embryos were transferred to the uterus of 2.5 dpc pseudopregnant *ICR* females (Charles River Laboratories Japan, Inc.) on or the next day of injection.

### Culture for derivation of stem cells

The culture medium for derivation of ES-like stem cells consists of Glasgow-modified eagles medium (GMEM, Sigma), 15% KnockOut Serum Replacement^®^ (KSR, Life Technologies), 1 non-essential amino acids (NEAA, Nakarai), 1 Sodium Pyruvate (Nakarai), 10^−4^M 2-mercaptoethanol (Nakarai), 1,000 U/ml of LIF and 10 mM ACTH (Kurabo on consignment). We confirmed the medium is optimal for the culture of conventional ES cells. The culture medium for derivation of TS-like stem cells consists of GMEM, 20% FCS, 1 NEAA, 1 Sodium Pyruvate, 10^−4^ M 2-mercaptoethanol, 25 ng/ml of recombinant mouse Fgf4 (Wako) and 1 mg/ml of heparin (Wako). We confirmed the medium is optimal for the culture of conventional TS cells. To derive stem cells, cell aggregates were isolated under a microscope and transferred into a well of 96-well plate with 100 ml of the culture medium and 1,000 feeder cells. Feeder cells were prepared by treatment of mouse embryonic fibroblasts prepared from day 14 C57BL6 embryos with Mitomycin C (Wako) for 3 hours.

## Acknowledgements

We would like to thank the assistance of the members of the Scientific Validity Examination Team, Dr. Hiroshi Kiyonari and Mr. Kenichi Inoue for chimera production and animal breeding, and Laboratory of Animal Resources and Genetic Engineering for animal housing. We also thank Mr. Douglas Sipp for critical discussion of this report. This examination was supported by the grant for Scientific Validity Examination by RIKEN President.

## References

Abe, T., Kiyonari, H., Shioi, G., et al. (2011) Establishment of conditional reporter mouse lines at ROSA26 locus for live cell imaging. Genesis 49, 579–590.

Bradley, A., Evans, M., Kaufman, M.H. & Robertson, E. (1984) Formation of germ-line chimaeras from embryo-derived teratocarcinoma cell lines. Nature 309, 255–256.

Evans, M.J. & Kaufman, M.H. (1981) Establishment in culture of pluripotential cells from mouse embryos. Nature 292, 154–156.

Gurdon, J.B. (1962) Adult frogs derived from the nuclei of single somatic cells. Dev Biol 4, 256–273.

Martin, G.R. (1981) Isolation of a pluripotent cell line from early mouse embryos cultured in medium conditioned by teratocarcinoma stem cells. Proc Natl Acad Sci U S A 78, 7634–7638.

Mizuno, N. & Kosaka, M. (2008) Novel variants of Oct-3/4 gene expressed in mouse somatic cells. J Biol Chem 283, 30997–31004.

Niwa, H., Toyooka, Y., Shimosato, D., et al. (2005) Interaction between Oct3/4 and Cdx2 determines trophectoderm differentiation. Cell 123, 917–929.

Obokata, H., Wakayama, T., Sasai, Y., et al. (2014a) Stimulus-triggered fate conversion of somatic cells into pluripotency. Nature 505, 641–647.

Obokata, H., Sasai, Y., Niwa, H., et al. (2014b) Bidirectional developmental potential in reprogrammed cells with acquired pluripotency. Nature 505, 676–680.

Obokata, H., Wakayama, T., Sasai, Y., et al. (2014c) Retraction: Stimulus-triggered fate conversion of somatic cells into pluripotency. Nature 511, 112.

Obokata, H., Sasai, Y., Niwa, H., et al. (2014d) Retraction: Bidirectional developmental potential in reprogrammed cells with acquired pluripotency. Nature 511, 112.

Ogawa, K., Matsui, H., Ohtsuka, S. & Niwa, H. (2004) A novel mechanism for regulating clonal propagation of mouse ES cells. Genes Cells 9, 471–477.

Ohbo, K., Yoshida, S., Ohmura, M., et al. (2003) Identification and characterization of stem cells in prepubertal spermatogenesis in mice. Dev Biol 258, 209–225.

Ohtsuka, S. & Niwa, H. (2015) The differential activation of intracellular signaling pathways confers the permissiveness of embryonic stem cell derivation from different mouse strains. Development 142, 431–437.

Okabe, M., Ikawa, M., Kominami, K., Nakanishi, T. & Nishimune, Y. (1997) ‘Green mice’ as a source of ubiquitous green cells. FEBS Lett 407, 313–319.

Okamoto, K., Okazawa, H., Okuda, A., Sakai, M., Muramatsu, M. & Hamada, H. (1990) A novel octamer binding transcription factor is differentially expressed in mouse embryonic cells. Cell 60, 461–472.

Postic, C., Shiota, M., Niswender, K.D., et al. (1999) Dual roles for glucokinase in glucose homeostasis as determined by liver and pancreatic beta cell-specific gene knock-outs using Cre recombinase. J Biol Chem 274, 305–315.

Smith, A.G., Heath, J.K., Donaldson, D.D., et al. (1988) Inhibition of pluripotential embryonic stem cell differentiation by purified polypeptides. Nature 336, 688–690.

Takahashi, K. & Yamanaka, S. (2006) Induction of pluripotent stem cells from mouse embryonic and adult fibroblast cultures by defined factors. Cell 126, 663–676.

Tanaka, S., Kunath, T., Hadjantonakis, A.K., Nagy, A. & Rossant, J. (1998) Promotion of trophoblast stem cell proliferation by FGF4. Science 282, 2072–2075.

Yeom, Y.I., Fuhrmann, G., Ovitt, C.E., et al. (1996) Germline regulatory element of Oct-4 specific for the totipotent cycle of embryonal cells. Development 122, 881–894.

